# Baculovirus actin-rearrangement-inducing factor ARIF-1 induces the formation of dynamic clusters of invadosome-like structures

**DOI:** 10.1101/2020.11.03.365502

**Authors:** Domokos I. Lauko, Taro Ohkawa, Sergio E. Mares, Matthew D. Welch

## Abstract

The baculovirus *Autographa californica* multiple nucleopolyhedrovirus (AcMNPV), a pathogen of lepidopteran insects, has a striking dependence on the host cell actin cytoskeleton. During the delayed-early stage of infection, AcMNPV was shown to induce the accumulation of actin at the cortex of infected cells. However, the dynamics and molecular mechanism of cortical actin assembly remained unknown. Here, we show that AcMNPV induces dynamic cortical clusters of dot-like actin structures that resemble clusters of invadosomes in mammalian cells. Furthermore, we find that the AcMNPV protein actin-rearrangement-inducing factor-1 (ARIF-1), which was previously shown to be necessary and sufficient for cortical actin assembly and efficient viral infection in insect hosts, is both necessary and sufficient for invadosome-like structure formation. We mapped the sequences within the C-terminal cytoplasmic region of ARIF-1 that are required for invadosome-like structure formation and identified individual tyrosine and proline residues that are required for organizing these structures. Additionally, we found that ARIF-1 and the invadosome-associated proteins cortactin and the Arp2/3 complex localize to invadosome-like structures, and Arp2/3 complex is required for their formation. These ARIF-1-induced invadosome-like structures may be important for the function of ARIF-1 in systemic virus spread.

## Introduction

Intracellular microbial pathogens are master manipulators of host cell biology and have evolved myriad strategies to hijack host cell machinery and repurpose host processes to facilitate infection. One such strategy is to hijack the host actin cytoskeleton, which can facilitate pathogen invasion, intracellular movement, and/or cell-cell spread. The baculovirus *Autographa californica* multiple nucleopolyhedrovirus (AcMNPV), an enveloped DNA virus that orally infects larval lepidopteran insects (caterpillars), is notable for manipulating the host actin cytoskeleton extensively throughout infection. Upon entry into the host cell cytoplasm, AcMNPV nucleocapsids undergo actin-based motility, using the viral P78/83 protein to activate host Arp2/3 complex to polymerize actin filaments (Goley et al., 2006; Ohkawa et al., 2010). Upon expression of early viral genes, actin filaments accumulate at the cortex of infected cells (Charlton and Volkman, 1991; Dreschers et al., 2001; Roncarati and Knebel-Mörsdorf, 1997). This accumulation dissipates during late viral gene expression as monomeric actin is imported into and polymerizes within the nucleus (Charlton and Volkman, 1993, 1991; Hepp et al., 2018; Ohkawa et al., 2002; Ohkawa and Volkman, 1999; Volkman et al., 1992). Newly assembled viral nucleocapsids also harness P78/83 and Arp2/3 complex to polymerize actin and are propelled to the nuclear periphery to facilitate nuclear egress (Ohkawa and Welch, 2018). As both actin polymerization in the nucleus and viral actin-based motility are required for successful viral infection (Goley et al., 2006; Hess et al., 1989; Ohkawa and Volkman, 1999; Volkman, 1988; Volkman et al., 1987; Volkman and Kasman, 2000), the ability of AcMNPV to hijack the host actin cytoskeleton is of critical importance.

A second AcMNPV protein that impacts the actin cytoskeleton is the actin-rearrangement-inducing factor-1 (ARIF-1) (Roncarati and Knebel-Mörsdorf, 1997). ARIF-1 is a delayed-early viral protein that was identified in a screen for AcMNPV genes that cause alterations in the actin cytoskeleton when expressed in insect cells, and was shown to be necessary and sufficient to induce the accumulation of actin filaments at the plasma membrane during early infection (Roncarati and Knebel-Mörsdorf, 1997). *arif-1* is conserved in alphabaculoviruses (Roncarati and Knebel-Mörsdorf, 1997), and ARIF-1 contains three predicted transmembrane domains and a C-terminal proline-rich cytoplasmic domain (Dreschers et al., 2001). Although ARIF-1 localizes to the plasma membrane in infected cells and is phosphorylated on tyrosine residues (Dreschers et al., 2001), the mechanism by which it induces actin polymerization is unknown. Interestingly, ARIF-1 is not important for viral replication in cultured cells (Dreschers et al., 2001; Kokusho et al., 2015; Taka et al., 2013). However, insects infected with an *arif-1* mutant *Bombyx mori* NPV (BmNPV; closely related to AcMNPV) experienced delays in infection of major organ systems and death, indicating that ARIF-1 accelerates systemic infection (Kokusho et al., 2015). Nevertheless, the mechanisms by which ARIF-1 functions in rearrangement of the host actin cytoskeleton in cells, and systemic infection in caterpillars, are still unknown.

During infection in caterpillars, AcMNPV must bypass barriers to systemic infection. AcMNPV infects through the oral route and establishes initial infection in midgut epithelial cells (Rohrmann, 2019). After replication, the virus spreads from the midgut to the tracheal system (Engelhard et al., 1994) and then to most major organ systems. However, the basal lamina (BL), a layer of extracellular matrix (ECM) that surrounds the midgut and other major organ systems, represents a barrier to virus spread (Passarelli, 2011), as gaps or pores in the BL are thought to be too small for viral particles to cross (Hess and Falcon, 1987; Reddy and Locke, 1990). Because AcMNPV is found in the caterpillar circulatory system only 30 min post-infection (Granados and Lawler, 1981), the virus must possess mechanisms to rapidly bypass the BL.

An unexplored mechanism for BL penetration is that of BL breakdown using actin-rich podosomes and invadopodia (collectively known as invadosomes). These are dot-like actin-containing structures of 0.5 to 2 μm in diameter (Marchisio, 1987; Marchisio et al., 1984; Nermut et al., 1991; Tarone et al., 1985) that may be organized into dynamic clusters shaped as rings or rosettes (Destaing et al., 2003; Kuo et al., 2018). Invadosomes are characterized by the presence of a dynamic actin core (Destaing et al., 2003) as well as the presence of the actin-associated proteins cortactin (Hiura et al., 1995; Kanner et al., 1990; Oser et al., 2009; Schuuring et al., 1993) and the Arp2/3 complex (Burns et al., 2001; Linder et al., 2000, 1999; Yamaguchi et al., 2005), as well as the scaffold protein Tks5 (Di Martino et al., 2014; Seals et al., 2005). Invadosomes are also sites of directed ECM degradation by matrix metalloproteases (Burgstaller and Gimona, 2005; Chen, 1989; Chen et al., 1985). Invadosomes are common in monocyte-derived cells such as osteoclasts that penetrate the ECM during migration or remodel the ECM (Marchisio, 1987; Marchisio et al., 1984), though they also occur in smooth muscle cells (Hai et al., 2002; Webb et al., 2005; Zhou et al., 2006) and endothelial cells (Moreau et al., 2003; Varon et al., 2006). Formation of invadosomes is also associated with aggressive cancer cell lines and enables them to remodel the ECM and undergo metastasis (reviewed in (Eddy et al., 2017; Paz et al., 2014). Vertebrate tumor virus infection can also induce invadosome formation. For example, transformation of fibroblasts with Rous sarcoma virus leads to expression of viral-Src (v-Src), a tyrosine kinase that can induce invadosome formation through activation of Tks5 (Chen, 1989; David-Pfeuty and Singer, 1980; Seals et al., 2005; Stylli et al., 2009; Tarone et al., 1985). Whether and how viruses that infect invertebrate animals induce invadopodia has remained uncertain.

Upon investigating AcMNPV-induced cortical actin rearrangements, we found that the virus promotes the formation of actin-containing structures in lepidopteran cells whose appearance and dynamics are similar to invadosome clusters in mammalian cells. Furthermore, we found that ARIF-1 is necessary and sufficient for formation of these invadosome-like structures. We mapped the regions of ARIF-1 and identified individual tyrosine and proline residues that are necessary for formation of clusters of invadosome-like structures. Finally, we observed that invadosome-like structures co-localize with ARIF-1, cortactin, and the Arp2/3 complex, and Arp2/3 complex activity is required for their formation and maintenance. Our findings suggest that ARIF-1 induces the formation of invadosome-like structures that may play a role in accelerating viral infection in insect hosts.

## Results

### AcMNPV infection induces the formation of invadosome-like structures

To investigate actin cytoskeleton rearrangements induced by AcMNPV during the early stage of infection, we transiently transfected *Spodoptera frugiperda* Sf21 cells with a plasmid expressing GFP-tagged actin (GFP-actin) and infected them with AcMNPV. As soon as 3 h post-infection (hpi), cells formed striking actin structures that appeared as small round clusters, circular rosettes, or elongated belts, ranging between 3 and 20 μm in size **(Fig 1A)**. TIRF microscopy revealed that these structures were basally located on the substrate-facing cell surface **(Fig 1A)**. We then quantified the percentage of cells with these actin structures over the course of viral infection. We found that actin structures began to form in cells at 3 to 4 hpi, were most prevalent between 4 and 8 hpi, and could be found at times as late as 32 hpi **(Fig 1B)**.

**Figure 1:**
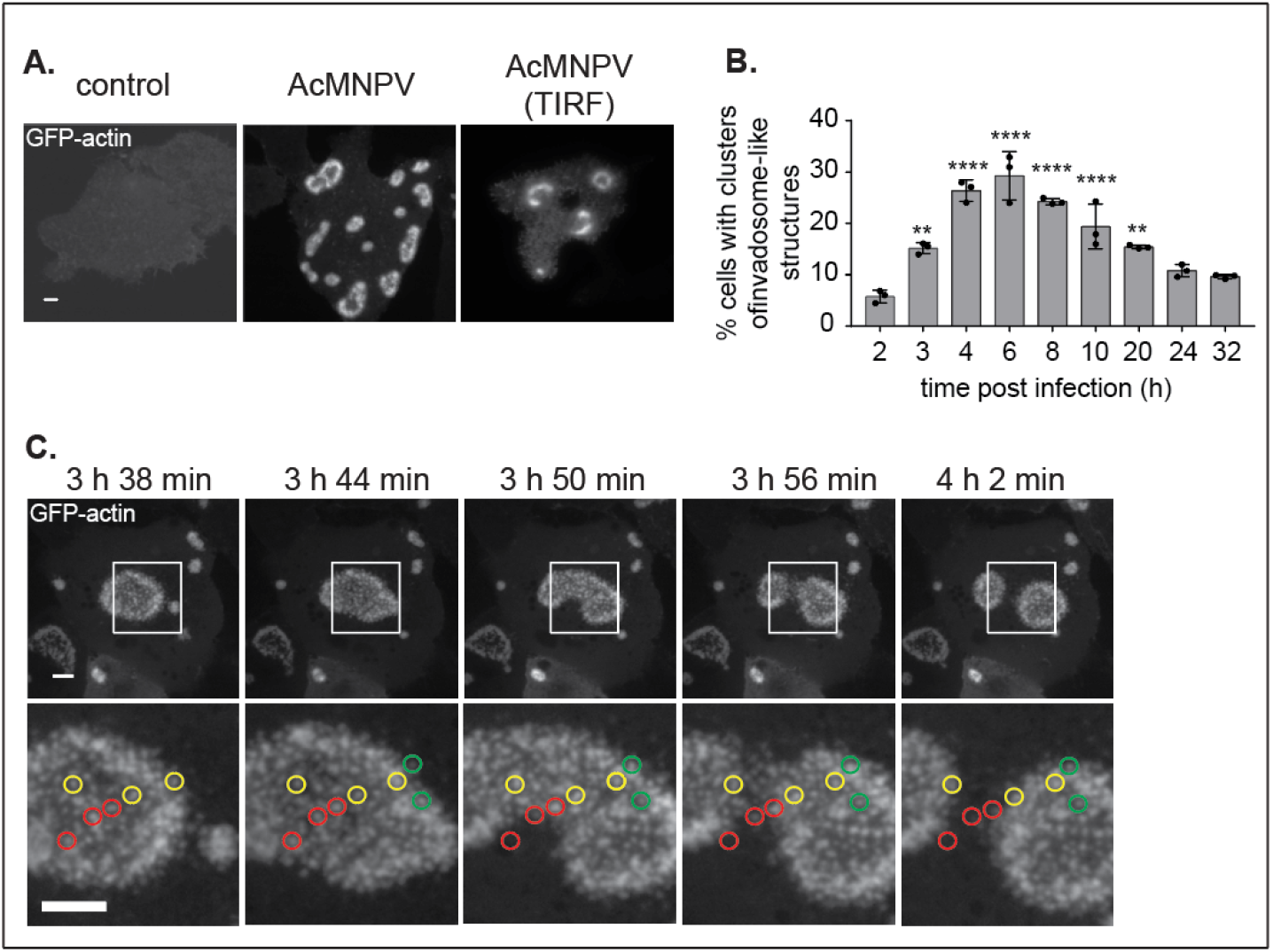
Dynamic clusters of invadosome-like structures form in Sf21 insect cells during early AcMNPV infection. **(A)** Confocal and TIRF images of Sf21 insect cells transiently transfected with GFP-actin and mock infected or infected with AcMNPV MOI of 10. Images were taken 4 to 7 hpi. Scale bars = 5 μm. **(B)** Clusters of invadosome-like actin structures in AcMNPV infected Sf21 cells were quantified from 2 to 32 hpi. Data are mean ± SD of n=3 biological replicates imaging approximately 5000 cells each. P-values were calculated with a one-way ANOVA with Tukey’s post-hoc test relative to the 2h timepoint and are indicated as follows: ** = p<0.005, **** = p<0.0001. **(C)** Confocal time-lapse images of Sf21 insect cells transiently transfected with GFP-actin and infected with AcMNPV. Inset images on the bottom row show red, yellow, and green circles enclosing stationary actin puncta that disappear, are maintained, or appear respectively. Scale bars = 5 μm. Time post-infection is indicated.

To further investigate the dynamics of these structures, we imaged live, infected Sf21 cells from 0 to 8 hpi. We found that the actin structures were highly dynamic in shape and position, could persist for more than 4.5 h, and could undergo fusion or fission events **(Fig 1C**, **Video S1)**. Interestingly, the larger actin structures were comprised of many smaller ~0.5 μm dot-like actin puncta **(Fig 1C**, **Video S2)**. Individual actin puncta remained stationary relative to the substrate, and the shape of the cluster changed through appearance or disappearance of individual actin puncta **(Fig 1C, Video S2)**. The appearance and behavior of the actin structures in AcMNPV-infected Sf21 cells were markedly similar to invadosome rings and rosettes found in some mammalian cell types, prompting us to designate them as clusters of invadosome-like structures.

To investigate the kinetics of actin polymerization-depolymerization in invadosome-like structures, we added latrunculin A (latA) to live infected cells and measured their persistence. Addition of latA caused GFP-actin signal in clusters of invadosome-like structures to rapidly fade with a half-life of ~7-8 min, so that none of the structures remained ~15 min after drug addition **(Video S3, Fig S1)**. The rapid disappearance of invadosome-like structures indicates that actin disassembly in these structures is relatively rapid, and that continuous actin polymerization is needed for their assembly and maintenance.

### ARIF-1 is necessary and sufficient for formation of invadosome-like structures

We next set out to define the viral gene(s) required for the formation of AcMNPV-induced invadosome-like structures. The AcMNPV *arif-1* gene was previously shown to be necessary and sufficient to induce accumulation of actin filaments at the cell periphery in TN368 insect cells (Roncarati and Knebel-Mörsdorf, 1997), suggesting that it may be responsible for inducing the formation of invadosome-like structures in Sf21 cells. To determine whether ARIF-1 plays a role in the formation of these structures, we constructed an *AcΔarif-1* virus in the AcMNPV WOBpos background that contained a deletion of 70% of the *arif-1* coding region (WOBpos is derived from the E2 strain of AcMNPV and its genome can be propagated as a bacmid in *Escherichia coli*). We also constructed an *AcΔarif-1 rescue* virus in which the *arif-1* gene and 500 bp flanking regions were inserted at the nearby *polyhedrin* locus in the *AcΔarif-1* viral genome. Sf21 cells infected with *AcΔarif-1* completely lacked clusters of invadosome-like structures **(Fig 2A, B)**. Formation of clusters of invadosome-like structures was fully restored in cells infected with the *AcΔarif-1*-*rescue* virus **(Fig 2A, B)**. This demonstrates that ARIF-1 is necessary for invadosome-like structure formation in infected Sf21 cells.

**Figure 2:**
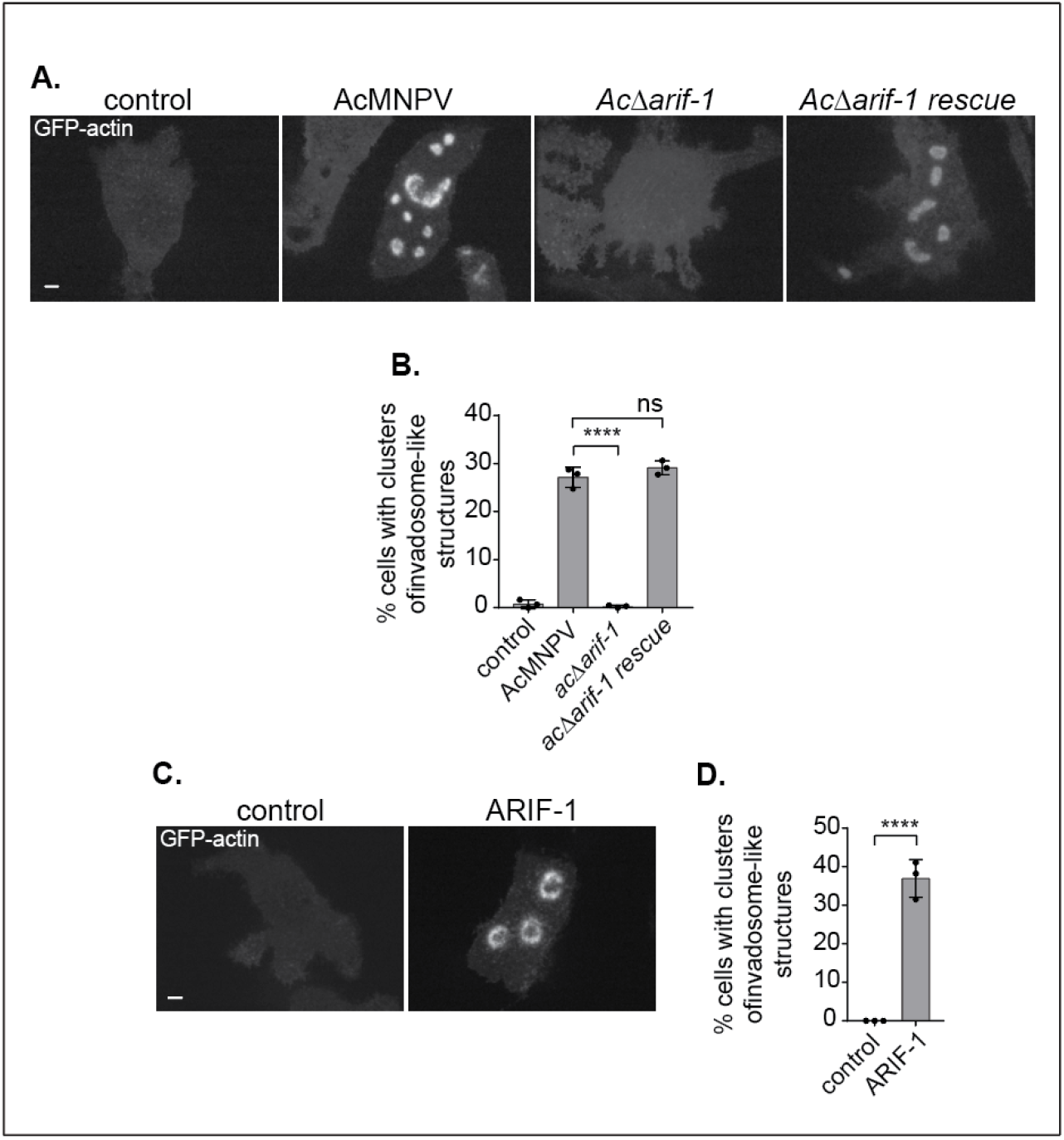
ARIF-1 is necessary and sufficient for formation of clusters of invadosome-like structures. **(A)** Confocal images of Sf21 cells transiently transfected with GFP-actin and infected with MOI of 10 of the indicated virus. Images were taken at 4 hpi and are representative of three biological replicates. Scale bars = 5 μm. **(B)** Clusters of invadosome-like structures in infected cells were quantified at 4 hpi. Data are mean ± SD of n=3 biological replicates of approximately 5000 cells each. P-values were calculated by one-way ANOVA with multiple comparisons and are indicated as follows: ns = not significant, **** = p<0.0001. **(C)** Images of Sf21 cells transiently expressing GFP-actin and ARIF-1. Images were taken 2 d post-transfection and are representative of three biological replicates. Scale bars = 5 μm. **(D)** Quantification of clusters of invadosome-like structures in cells transfected with GFP-actin and ARIF-1. Structures were quantified by eye 2 d post-transfection. Data are mean ± SD of n=3 biological replicates of 60 cells each treatment. The p-value was calculated by an unpaired t-test; **** = p = 0.0002.

To determine if ARIF-1 is sufficient for invadosome-like structure formation, we transiently transfected Sf21 cells with plasmid pACT-*arif-1*, which included *arif-1* under the control of the *B. mori* actin promoter. Transfected Sf21 cells formed clusters of invadosome-like structures **(Fig 2C, D)** that maintained a similar variety of shapes to those in infected cells (small and round, circular rosettes, or elongated belts). The dynamic behavior of these structures was also similar to that in infected cells **(Video S4)**. Overall, these data indicate that expression of *arif-1* alone is sufficient for the formation of clusters of invadosome-like structures in Sf21 cells.

To verify that the timing of ARIF-1 expression and that of invadosome-like structure formation were consistent with one another, we raised a polyclonal ARIF-1 antibody and probed lysates of cells infected with AcMNPV, *AcΔarif-1*, and *AcΔarif-1*-*rescue* viruses over a time course of infection **(Fig S2)**. ARIF-1 was expressed in cells infected with wild-type and *AcΔarif-1 rescue* viruses, but absent from cells infected with *AcΔarif-1*. Furthermore, the timing of ARIF-1 expression was consistent with the timing of invadosome-like structure formation and disappearance **(Fig 1B, Fig S2)**. This confirms that invadosome-like structure formation is correlated with ARIF-1 expression.

### ARIF-1 is localized to clusters of invadosome-like structures

ARIF-1 was previously reported to exhibit general localization to the plasma membrane in TN368 cells (Dreschers et al., 2001). To determine if ARIF-1 concentrates in invadosome-like structures, we investigated the localization of ARIF-1 in Sf21 cells. In cells transiently transfected with GFP-tagged ARIF-1 (GFP-ARIF-1), GFP-ARIF-1 localized to the plasma membrane, with a concentration at clusters of invadosome-like structures **(Fig 3)**. In some instances, the localization appeared uniform throughout the cluster, and in others it appeared to be more prominent at the periphery of the cluster **(Fig 3)**. Thus, ARIF-1 is enriched at clusters of invadosome-like structures.

**Figure 3:**
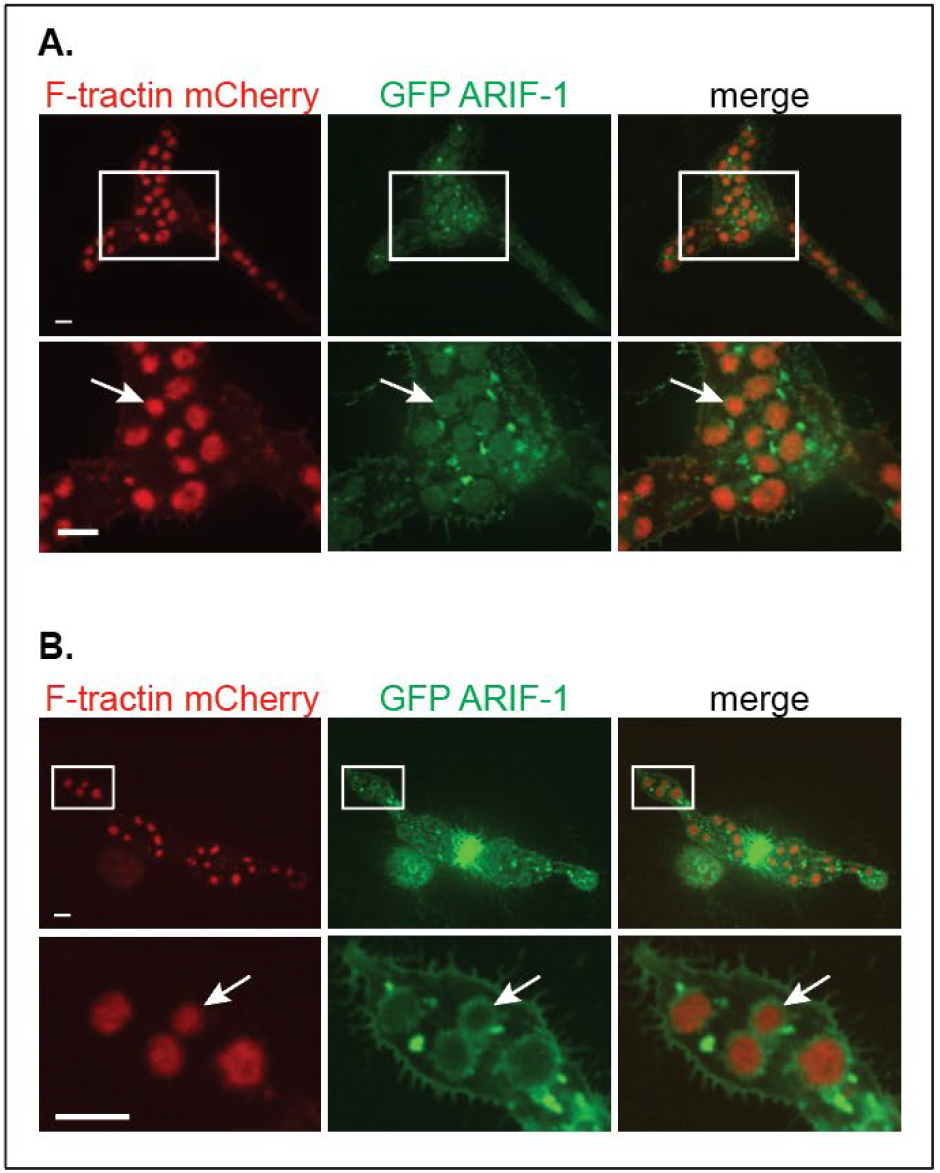
ARIF-1 co-localizes with clusters of invadosome-like structures. **(A)** Confocal images of Sf21 cells transiently expressing F-tractin mCherry and GFP-tagged ARIF-1. Images were taken 2 d post-transfection. Insets show areas of co-localization of mCherry and GFP signal, indicated by arrows. Scale bars = 5 μm. **(B)** Confocal images of Sf21 cells transiently expressing F-tractin mCherry and GFP-tagged ARIF-1. Images were taken 2 d post-transfection. Insets show areas of mCherry signal surrounded by GFP signal, as indicated by arrows. Scale bars = 5 μm.

### The ARIF-1 C-terminal region is necessary and sufficient for formation of clusters of invadosome-like structures

Prior structural predictions and our own analyses suggested that ARIF-1 contains three N-terminal transmembrane domains and a ~200 aa C-terminal region that extends into the cytoplasm (Dreschers et al., 2001) **(Fig. 4A)**. We first sought to determine which parts of the ARIF-1 C-terminal region are necessary for formation of clusters of invadosome-like structures. We constructed a series of C-terminal truncations of ARIF-1 and quantified formation of clusters of invadosome-like structures in transiently transfected Sf21 cells **(Fig 4B, Fig S3A)**. Although expression of ARIF-1(1-401) (containing C-terminal residues up through amino-acid 401), ARIF-1(1-398), ARIF-1(1-378), and ARIF-1(1-371) caused a reduced percentage of cells with clusters of invadosome-like structures when compared with expression of the full-length protein ARIF-1(1-417), these clusters still formed. However, no such structures formed in cells transfected with ARIF-1(1-274). This indicates that the ARIF-1 C-terminus between residues 274-371 is necessary for formation of clusters of invadosome-like structures.

**Figure 4:**
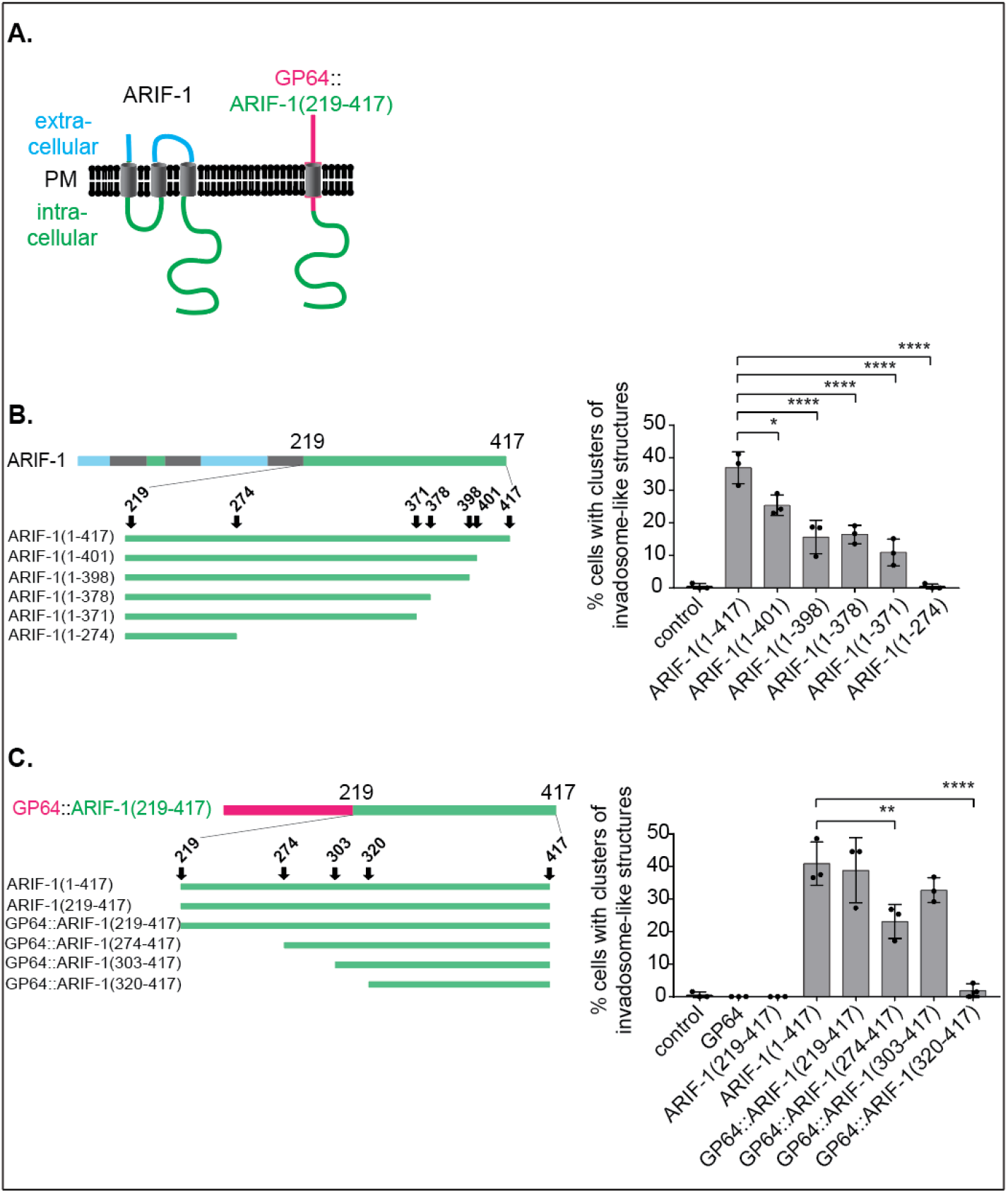
ARIF-1 residues 303-371 are necessary for formation of clusters of invadosome-like structures. **(A)** Left: predicted ARIF-1 structure with three transmembrane domains and a cytoplasmic C-terminal region (amino acids 219-417). Right: AcMNPV transmembrane protein GP64 (red) fusion to ARIF-1 C-terminal region. **(B)** Left: visual representation of ARIF-1 C-terminal truncations. Right: quantification of clusters of invadosome-like structures in Sf21 cells transiently expressing GFP-actin and truncated ARIF-1 2 d post-transfection. Data are mean ± SD of n=3 biological replicates of 60 cells each. P-values were calculated with a one-way ANOVA with multiple comparisons, comparing each treatment to ARIF-1 (1-417) and are indicated as follows: * = p=0.0167, **** = p<0.0001. **(C)** Left: visual representation of AcMNPV transmembrane protein GP64 (red) fused to ARIF-1(219-417) and N-terminal truncations. Right: quantification of clusters of invadosome-like structures in Sf21 cells transiently expressing GFP-actin and the indicated construct 2 d post-transfection. Data are mean ± SD of n=3 biological replicates of 60 cells each. P-values were calculated with a one-way ANOVA with multiple comparisons, comparing each treatment to ARIF-1 (1-417) and are indicated as follows: ** = p=0.0075, **** = p<0.0001.

To determine the contributions of the ARIF-1 N-terminal and transmembrane regions, we transfected cells with a plasmid that expressed ARIF-1(219-417) missing N-terminal amino acids 1-218 that encode for the predicted transmembrane domains and cytoplasmic loop **(Fig 4C)**. Sf21 cells transiently transfected with ARIF-1(219-417) did not form invadosome-like structures **(Fig 4C, Fig S3B)**, indicating that the transmembrane domains are required for ARIF-1 function. Next, to test whether membrane targeting of the C-terminus is sufficient for formation of clusters of invadosome-like structures, we expressed a variant of ARIF-1 in which the ARIF-1 C-terminus was fused to the unrelated AcMNPV transmembrane protein GP64 **(Fig 4A, right)**. Surprisingly, a similar percentage of cells expressing GP64::ARIF-1(219-417) formed clusters of invadosome-like structures compared with cells expressing full-length ARIF-1(1-417) **(Fig 4C, Fig S3B, Video S5)**. This indicates that the membrane-targeted ARIF-1 C-terminus from residues 219-417 is sufficient for formation of clusters of invadosome-like structures, and that the AIRF-1 N-terminal cytoplasmic loop and transmembrane regions function to anchor the ARIF-1 C-terminal region to the plasma membrane.

To further narrow down which regions of the ARIF-1 C-terminus are necessary for formation of clusters of invadosome-like structures, we constructed a series of N-terminal truncations to GP64::ARIF-1(219-417) and quantified invadosome-like structure formation in transiently transfected Sf21 cells. Cells expressing GP64::ARIF-1(274-417) and GP64::ARIF-1(303-417) had clusters of invadosome-like structures, whereas these structures were completely absent in cells expressing GP64::Arif-1(320-417) **(Fig 4C, Fig S3B)**. Altogether, the data from expression of various truncation derivatives indicate that the ARIF-1 C-terminus between residues 303-371 is necessary for formation of clusters of invadosome-like structures.

### ARIF-1 residues Y332F and P335A are important for formation of clusters of invadosome-like structures

We next sought to identify individual residues in the ARIF-1 C-terminal region that may be important for the formation of clusters of invadosome-like structures. ARIF-1 is tyrosine-phosphorylated during infection, though which tyrosine residues are phosphorylated is unknown (Dreschers et al., 2001). The ARIF-1 C-terminus also contains several stretches rich in proline residues **(Fig 5A, left)**. We sought to assess the importance of individual tyrosine and proline residues by mutating tyrosine to phenylalanine and proline to alanine **(Fig 5A)**. We then quantified clusters of invadosome-like structures in Sf21 cells transiently expressing ARIF-1 mutants **(Fig 5B, C)**. While most mutations did not significantly affect the formation of clusters of invadosome-like structures, cells transiently transfected with ARIF-1(Y332F) and ARIF-1(P335F) mutations had no clusters, but formed invadosome-like structures uniformly dispersed across the basal cell surface **(Fig 5D, Fig S4A, B, Video S6, Video S7)**. To verify that differences in formation of clusters of invadosome-like structures in ARIF-1 tyrosine and proline point mutants were not due to differences in expression levels of the ARIF-1 point mutant, we probed cell lysates using the ARIF-1 antibody **(Fig S5A, B)**. ARIF-1 was detected in cells transfected with ARIF-1 Y332F and P335A mutations, demonstrating that the lack of clusters of invadosome-like structures was not due to decreased ARIF-1 expression. Our results are consistent with truncation analyses, which pointed to key residues between 303-371, and suggest that adjacent residues Y332 and P335 facilitate the formation of clusters of invadosome-like structures.

**Figure 5.**
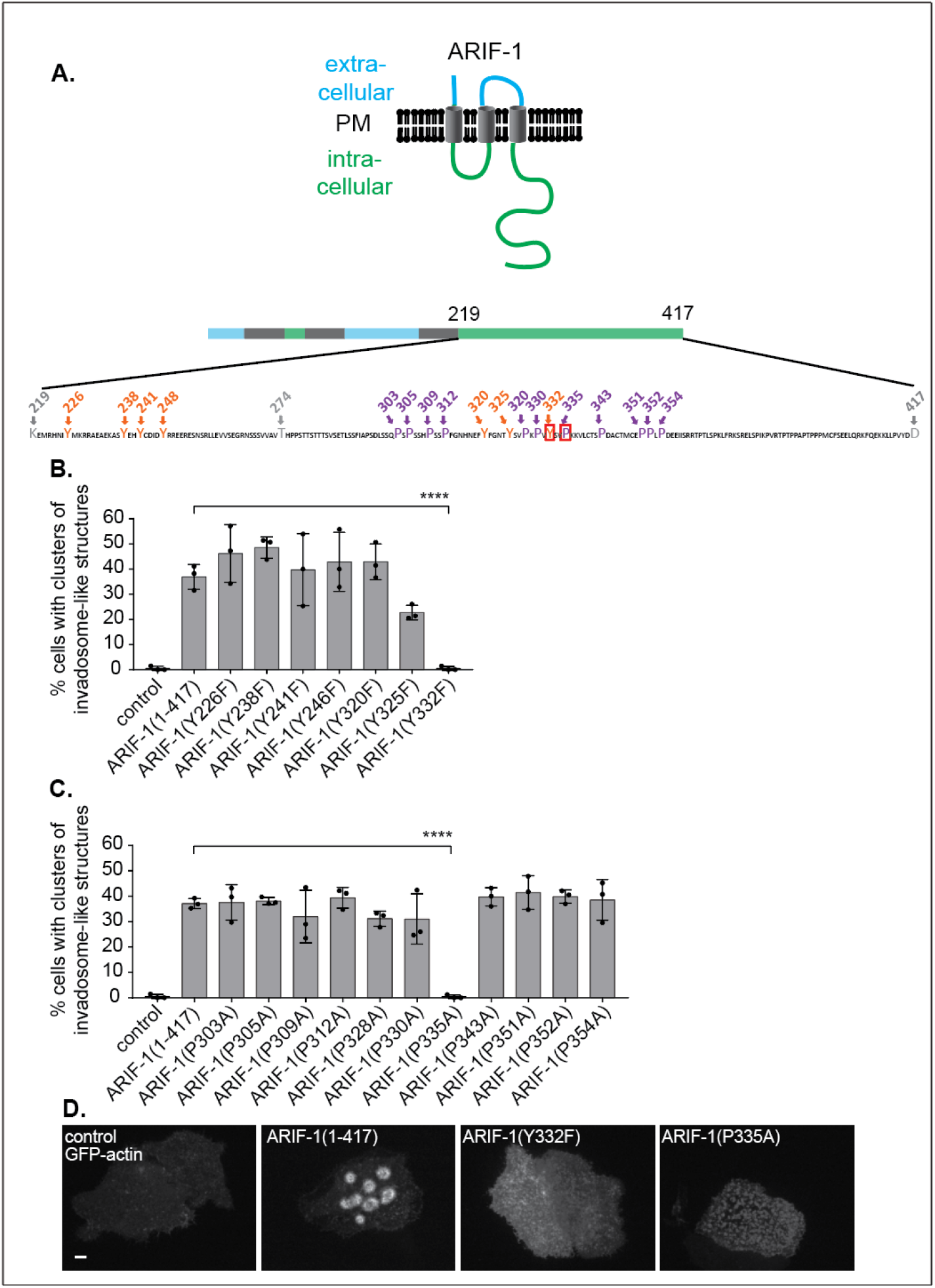
ARIF-1 residues Y332F and P335A are important for formation of clusters of invadosome-like structures. **(A)** Tyrosine (Y, orange) and proline (P, purple) residues are indicated on the C-terminal region of ARIF-1. **(B)** Clusters of invadosome-like structures in cells transiently transfected with GFP-actin and the indicated tyrosine to phenylalanine point mutations were quantified 2 d post-transfection. Data are mean ± SD of n=3 biological replicates of 60 cells each. P-values were calculated with a one-way ANOVA with multiple comparisons, comparing each treatment to ARIF-1 (1-417) and are indicated as follows **** = p<0.0001. **(C)** Clusters of invadosome-like structures in cells transiently transfected with GFP-actin and the indicated proline to alanine point mutations were quantified 2 d post-transfection. Data are mean ± SD of n=3 biological replicates of 60 cells each. P-values were calculated with a one-way ANOVA with multiple comparisons, comparing each treatment to ARIF-1 (1-417) and are indicated as follows: **** = p<0.0001. **(D)** Confocal images of Sf21 cells transiently expressing GFP-actin and the indicated ARIF-1 mutation. Images are representative of three biological replicates. Scale bars = 5 μm.

### Cortactin and the Arp2/3 complex play a role in the formation and maintenance of invadosome-like structures

The actin core of podosomes in mammalian cells includes cortactin (Hiura et al., 1995; Schuuring et al., 1993) and the Arp2/3 complex (Linder et al., 2000, 1999; Mizutani et al., 2002). To determine if invadosome-like structures in lepidopteran cells have a similar protein composition, we investigated whether cortactin and the Arp2/3 complex colocalize with these structures. GFP-tagged *S. frugiperda* cortactin (GFP-cortactin) was expressed in Sf21 cells by transient transfection, and cells were subsequently infected with AcMNPV WOBpos and imaged at 4 hpi. GFP-cortactin co-localized with actin in invadosome-like structures **(Fig 6A)**. To determine if the Arp2/3 complex localizes to clusters of invadosome-like structures, we expressed GFP-tagged Arp2/3 complex subunit ARPC3 (GFP-ARPC3) in Sf21 cells by transient transfection, and similarly infected these cells with AcMNPV and imaged at 4 hpi. GFP-ARPC3 also co-localized with clusters of invadosome-like structures **(Fig 6B)**. Thus, both cortactin and the Arp2/3 complex are present at these areas of dynamic actin polymerization.

**Figure 6:**
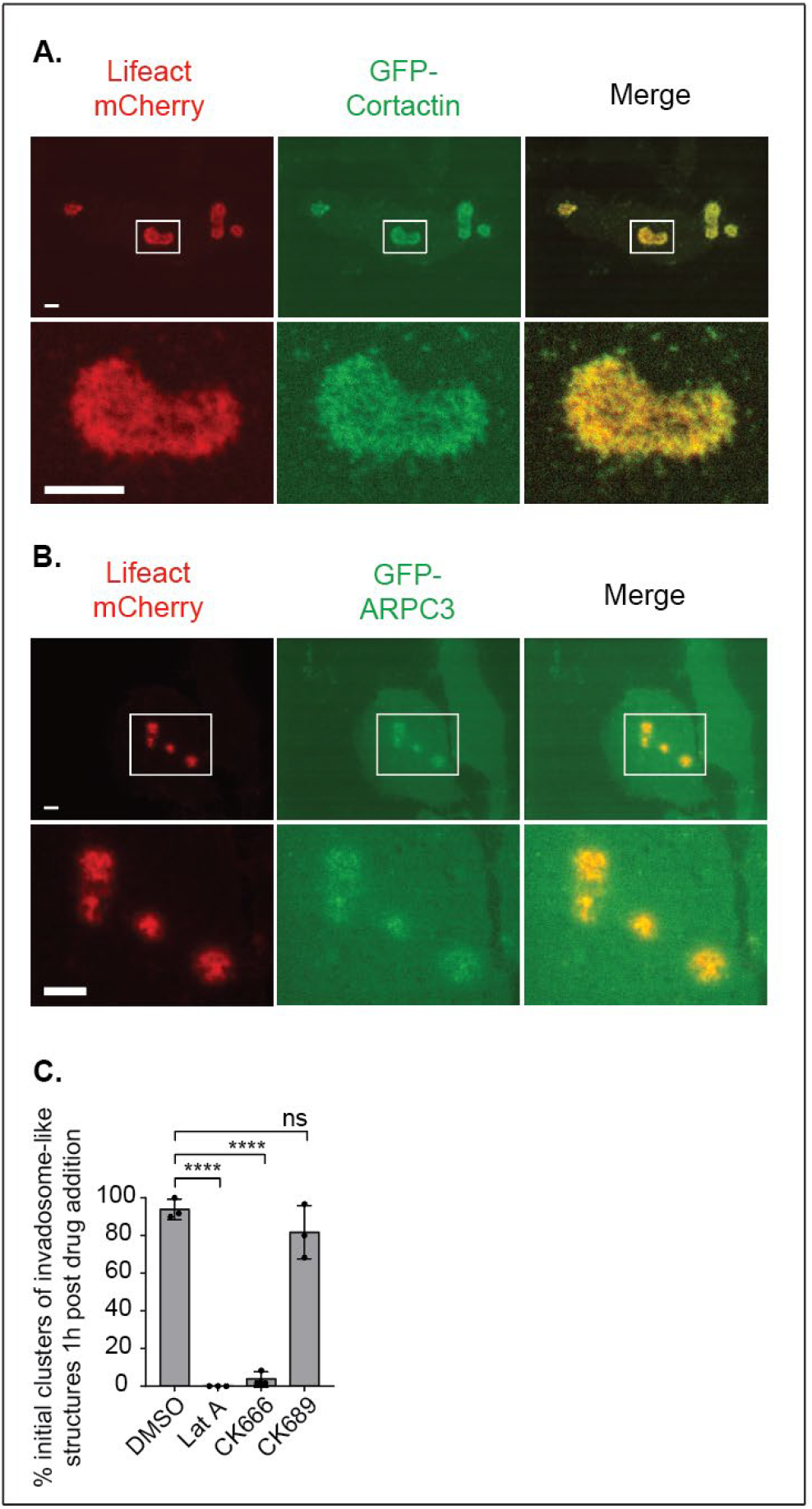
Cortactin and the Arp2/3 complex are present at clusters of invadosome-like structures, and Arp2/3 complex is required for their formation and maintenance. (A) Confocal images of Sf21 cells transiently expressing Lifeact mCherry and GFP-tagged *S.frugiperda* cortactin. Images were taken 2 d post-transfection and are representative of three biological replicates. Scale bars = 5 μm. (B) Confocal images of Sf21 cells transiently expressing Lifeact mCherry and GFP-tagged Arp2/3 complex subunit ARPC3. Images were taken 2 d post-transfection and are representative of three biological replicates. Scale bars = 5 μm. (C) Clusters of invadosome-like structures were quantified 1 h after the indicated drugs were added to Sf21 cells infected with AcMNPV MOI of 10 at 4 hpi. Data are mean ± SD of n=3 biological replicates of 40 invadosome-like structures. P-values were calculated with a one-way ANOVA with multiple comparisons, comparing each treatment to ARIF-1 (1-417) and are indicated as follows: ns = not significant, **** = p<0.0001.

To determine if the host Arp2/3 complex plays a role in formation and maintenance of invadosome-like structures, we treated infected cells with Arp2/3 complex inhibitor CK666, or with the inactive control drug CK869. Clusters of invadosome-like structures were virtually eliminated 1 h post-treatment with Arp2/3 inhibitor CK666, but no significant effect was observed upon treatment with CK869 **(Fig 6C, Video S7, S8, S9)**. Thus, host Arp2/3 complex function is required for formation and maintenance of these structures.

## Discussion

Here, we describe the formation of dynamic actin structures in AcMNPV-infected lepidopteran insect cells that coalesce into clusters, rosettes, and rings, resembling invadosome clusters in mammalian cells. We further show that the AcMNPV protein ARIF-1 is necessary and sufficient for formation of these invadosome-like structures. We identify regions of ARIF-1 and individual proline and tyrosine residues critical for their formation. Lastly, we verify that ARIF-1, cortactin, and the Arp2/3 complex localize with clusters of invadosome-like structures, and that Arp2/3 complex function is important for their maintenance. Our results suggest that ARIF-1 induces the formation of invadosome-like structures in lepidopteran cells, and that these structures may facilitate systemic AcMNPV spread in hosts.

Our results add to previous observations by describing the formation of AcMNPV-induced and ARIF-1-dependent invadosome-like structures. Previously, it was noted that during early-stage AcMNPV infection, actin accumulated evenly around the periphery of TN368 and BmN cells (Kokusho et al., 2015; Roncarati and Knebel-Mörsdorf, 1997), and accumulated in “ventral aggregates” on the basal cell surface in Sf21 cells (Charlton and Volkman, 1991). However, the finer organization and dynamics of this peripheral actin was not described. Using live cell imaging, we observed that ARIF-1 induces the formation of actin puncta that cluster together into dynamic clumps, rosettes, or belts, and that these can persist for hours and change shape and position. Though we have not observed clusters of invadosome-like structures in TN368 cells, they are present in infected Sf9 (data not shown), Sf21 (our work; Charlton and Volkman, 1991), and BmN (data not shown) cell lines, indicating that they are not restricted to a single cell line or even species.

ARIF-1-induced actin structures in Sf21 cells are similar to podosomes and invadopodia in appearance and dynamics. In mammalian osteoclasts, many stationary dot-like podosomes organize into clusters that merge to form one large ring around the cell periphery (Destaing et al., 2003; reviewed in Luxenburg et al., 2007). The shape of these ring structures is determined by selective activation and inactivation of stationary podosomes (Destaing et al., 2003), in the same way that individual pixels on a screen turn on and off to create a moving image. We have observed a similar phenomenon in clusters of invadosome-like structures in AcMNPV-infected or *arif-1*-transfected Sf21 cells. However, instead of full podosome rings as seen in osteoclasts, these structures more closely resemble invadopodia rosettes in v-Src-transformed fibroblast cells (Tarone et al., 1985). Intriguingly, invadopodia rosettes have also been described as dynamically changing shape (Stickel and Wang, 1987), fusing together (Kuo et al., 2018), or splitting in two to form new invadopodia rosettes (Kuo et al., 2018), all of which we observed in clusters of invadosome-like structures in lepidopteran cells. Individual osteoclast podosomes persist for 2 min on average, with actin turnover on the order of 1 min (Destaing et al., 2003). Meanwhile, podosome clusters in these cells persist for several hours (Destaing et al., 2003). In lepidopteran cells, our results suggest that actin in invadosome-like structures has a half-life of approximately 7 min and clusters persist for hours.

ARIF-1-induced actin structures also have similar protein composition compared with podosomes and invadopodia. We confirmed that ARIF-1 localizes to the plasma membrane, concentrating around the clusters of invadosome-like structures. Furthermore, actin, cortactin, and the Arp2/3 complex localize to invadosome-like structures themselves. These are also critical components of mammalian podosomes and invadopodia (Burns et al., 2001; Destaing et al., 2003; Hiura et al., 1995; Kanner et al., 1990; Linder et al., 1999, 2000; Oser et al., 2009; Schuuring et al., 1993; Yamaguchi et al., 2005), implying a parallel between the invadosome-like structures in mammalian and lepidopteran cells. Interestingly, the scaffolding protein tyrosine kinase with five SH3 domains (Tks5), which is a distinct marker of podosomes and invadopodia in mammalian cell types (Burger et al., 2014; Di Martino et al., 2014; Seals et al., 2005; Stylli et al., 2009), lacks a clear ortholog in *S. frugiperda*. This suggests the possibility that ARIF-1 itself may be acting as a scaffolding protein, playing a role similar to mammalian Tks5 in invadosome assembly.

We identified sequences in the ARIF-1 C-terminal region as well as specific residues that are required for forming clusters of invadosome-like structures. While the ARIF-1 C-terminus from amino acids 303-371 is required for formation of invadosome-like structures, the residues Y332 and P335 are required for cluster formation. In cells expressing ARIF-1 with Y332F and P335A point mutations, invadosome-like structures were distributed across the entire basal plasma membrane rather than in defined clusters. That a phosphoablative tyrosine to phenylalanine mutation at residue 332 causes this phenotype suggests that Y332 may be phosphorylated. Phosphorylated tyrosine residues are key components of binding sites for Src homology 2 (SH2) domains (Buday et al., 2002; Filippakopoulos et al., 2009), which are common in proteins such as Grb-2 and Nck-1 that regulate actin cytoskeleton activity. A truncated ARIF-1 (ARIF-1(1-255)) is also tyrosine-phosphorylated during infection (Dreschers et al., 2001), indicating that additional ARIF-1 tyrosine residues are likely phosphorylated. Increased ARIF-1 phosphorylation as infection progresses was speculated to be correlated with disappearance of the ARIF-1-induced peripheral actin in TN368 cells (Dreschers et al., 2001).

Thus, ARIF-1 tyrosine phosphorylation at Y332 may play a role during early AcMNPV infection, possibly by generating a binding site for cellular or viral proteins that regulate actin polymerization in invadosome-like structures, leading to formation of organized clusters.

Our findings describe an ARIF-1-induced invadosome-like structure in insect cells. At the level of the host caterpillar, ARIF-1 has also been shown to play a role in viral spread (Kokusho et al., 2015). One explanation that could link the cellular-level and organismal-level functions of ARIF-1 is that ARIF-1-induced invadosome-like structures might mediate degradation of the underlying ECM and degrade barriers to viral infection, such as the insect BL. While we have not yet confirmed that ECM degradation underneath clusters of invadosome-like structures occurs, this hypothesis may inform current models of how other viruses, including human arbovirus pathogens, cross the BL in their insect hosts. For example, Eastern equine encephalitis virus disrupts the midgut BL of infected mosquitos (Weaver et al., 1988), though the mechanism of this disruption is unknown. While podosome formation has not previously been described in lepidopteran cell lines, invadosome-like structures have been described playing a role in myoblast fusion in *Drosophila*, though these structures were not organized into larger clusters (Haralalka et al., 2011; Onel and Renkawitz-Pohl, 2009; Sens et al., 2010). Such insect invadosome-like structures may hint at the existence of an ancestral podosome or invadopodium activation pathway retained by both mammalian and invertebrate animals that may be taken advantage of by viral pathogens. Thus, further understanding of the molecular mechanisms of podosome and invadopodia formation in both mammalian and insect cell models may uncover conserved roles in infection and other diseases.

## Materials and Methods

### Cell lines and viruses

*Spodoptera frugiperda* ovarian-derived Sf9 cells were maintained in suspension culture in ESF 921 media (Expression Systems, Davis, CA) and in adherent culture in Grace’s media (Gemini Bio-Products, West Sacramento, CA) with 10% fetal bovine serum (FBS; Gemini Bio-Products) on T-25 tissue culture flasks at 28°C. *S. frugiperda* ovarian-derived Sf21 cells were maintained in suspension culture in Grace’s media (Gemini Bio-Products) with 10% FBS (Gemini Bio-Products) and 0.1% Pluronic F-68 (Gibco from Thermo Fisher Scientific, Waltham, MA) at 28°C, and in adherent culture on T-25 tissue culture flasks in Grace’s media (Gemini Bio-Products; Expression systems) with 10% FBS (Gemini Bio-Products) at 28°C. *Bombyx mori* ovarian-derived BmN cells were a generous gift from Kostas Iatrou (Institute of Biosciences & Applications, National Centre for Scientific Research, Greece), through Don Jarvis (University of Wyoming). These cells were maintained in adherent culture on T-25 tissue culture flasks in TC-100 media (Gibco from Thermo Fisher Scientific) with 10% FBS (Gemini Bio-Products) at 28°C. MDA-MB-231 human breast adenocarcinoma cells were maintained in adherent culture on T-75 tissue culture flasks in Dubelco’s modified eagle medium (Gibco from Thermo Fisher Scientific) supplemented with 10% FBS (Atlas Biologicals, Fort Collins, CO) that was heat-inactivated at 65°C for 20 min. AcMNPV WOBpos virus (Goley et al., 2006), derived from AcMNPV E2, was used as the wild-type virus.

### Generation of recombinant viruses

To generate AcMNPV lacking a functional *arif-1* gene (*AcΔarif-1*), we constructed a transfer vector by subcloning a SalI/XhoI fragment of AcMNPV viral genomic fragment EcoRI A, which contains *arif-1,* into the XhoI site of pBluescript II SK+ (Addgene, Watertown, MA) to create the plasmid pEcoRI_ASalxh.pBSKS.rev. This plasmid was digested with MluI (New England Biolabs, Ipswich, MA), removing 74% of the *arif-1* coding region, which was replaced with a subcloned 5.4 kb fragment of MluI-digested pBlue-Tet, containing *lacZ* and tetracycline-resistance (*tetR*) genes (Goley et al., 2006; Ohkawa et al., 2005) to help in the selection of recombinant bacmids. The transfer vector was linearized by digestion with SmaI and ApaI (New England Biolabs) and purified by agarose gel electrophoresis. 30 fmol of DNA was co-electroporated with 0.2 μg of WOBpos bacmid DNA (containing the kanamycin-resistance (*kanR*) gene) into BW251143/pKD46 *E.coli* (Datsenko and Wanner, 2000), which expresses an arabinose-inducible recombinase on a plasmid with temperature-sensitive replication (Goley et al., 2006). Recombinant bacmids were selected by plating on Luria-Bertani (LB) agar plates with 50 μg/ml kanamycin (Gibco from Thermo Fisher Scientific) and 10 μg/ml tetracycline (MilliporeSigma, St. Louis, MO).

To generate an *AcΔarif-1* rescue virus (*AcΔarif-1 rescue*), we introduced the *arif-1* gene into the polyhedrin locus of the *AcΔarif-1* bacmid. To do this, we generated the transfer plasmid pARIF-1-Rescue-2 by PCR, amplifying a 2.2 kb fragment including *arif-1* and 500 bp 5’ and 3’ flanking sequences from pEcoRI_ASalxh.pBSKS.rev (Table 1), and inserted it using Gibson assembly (New England Biolabs; Gibson *et al*, 2009) into pWOBCAT (Ohkawa et al., 2010) (Table 1) amplified and linearized by PCR, inserting it upstream of a chloramphenicol resistance (*cat)* gene to help in the selection of recombinant bacmids. The resulting plasmid was digested with NotI and KasI (New England Biolabs) to remove a truncated *kanR* gene upstream of *arif-1*, which was replaced by ligating (Takara Bio USA) a 2 kb PCR-amplified fragment from pWOBpos2 (Goley et al., 2006) (Table 1) including the AcMNPV mini-F replicon and a full *kanR* gene in place of the truncated *kanR* gene. This plasmid transfer vector, pARIF-1-Rescue-2, was linearized through KpnI digestion (New England Biolabs) and purified by agarose gel electrophoresis. 30 fmol of DNA was electroporated with 0.2 μg of *AcΔarif-1* bacmid DNA into BW251143/pKD46 *E.coli* as described above, and bacteria were plated on LB agar plates with 25 μg/ml chloramphenicol and 10 μg/ml tetracycline (MilliporeSigma).

In all cases, positive colonies were grown in 2x YT media (MilliporeSigma) with 25 μg/ml kanamycin (Gibco from Thermo Fisher Scientific) for 18 hours at 37°C. Bacmid DNA was extracted and transfected into Sf9 cells using TransIT-Insect Transfection reagent (Mirius Bio, Madison, WI). Resulting virus was amplified by passaging in Sf9 cells, and correct homologous recombination verified through restriction enzyme digestion of viral DNA. PCR and sequencing of viral DNA was also used to confirm the presence of each desired genome modification.

### Plasmid construction for expression of wild type and mutant ARIF-1

To express full-length ARIF-1 and ARIF-1 C-terminal truncations, we used PCR to amplify the following AcMNPV *arif-1* regions from pEcoRI_ASalxh.pBSKS.rev (listed as amino acid numbers): ARIF-1(1-417), ARIF-1(1-219), ARIF-1(1-274), ARIF-1(1-371), ARIF-1(1-378), ARIF-1(1-398), and ARIF-1(1-401) along with C-terminal TAG stop codons (Table 1). Fragments were purified by agarose gel electrophoresis and individually subcloned into BamHI/NotI-digested pBluescript II KS+ (Addgene). Colonies positive for the plasmid were selected on LB agar with 100 μg/ml ampicillin (Gibco from Thermo Fisher Scientific), 100 μM 5-bromo-4-chloro-3-indolyl-β-D-galactopyranoside (X-Gal) and 100 μM Isopropyl β-d-1-thiogalactopyranoside (IPTG). Next, these plasmids were digested with BamHI and NotI (New England Biolabs) and subcloned into BamHI/NotI-digested pACT (Ohkawa et al., 2002). The resulting plasmids have a *B. mori* actin promoter driving expression of each ARIF-1 truncation.

To generate ARIF-1 C-terminally tagged with eGFP-FLAG as well as GlyGlyGlyGlySer-eGFP-FLAG (with a N-terminal linker), we amplified two DNA fragments with PCR and used Gibson assembly to subclone these into the BamHI site of pBluescript II KS+ (Addgene). The first fragment, containing *arif-1,* was amplified from pEcoRI_ASalxh.pBSKS.rev with reverse primers incorporating or not incorporating a C-terminal GlyGlyGlyGlySer linker (Table 1). The second fragment, containing *eGFP,* was amplified from pEGFPN1 (Takara Bio USA) with the reverse primer incorporating a FLAG tag (Table 1). Colonies positive for the assembled plasmid were selected on LB agar with 100 μg/ml ampicillin and 0.1mM X-Gal and IPTG. We then PCR-amplified the ARIF-1-eGFP-FLAG or ARIF-1-GlyGlyGlyGlySer-eGFP-FLAG sequence and used Gibson assembly to subclone it into the NotI site of pACT (Table 1).

To generate fusions of ARIF-1 and its N-terminal truncations to AcMNPV GP64, we PCR-amplified 2 fragments and assembled them into the NotI site of pACT. The first fragment was *gp64* from the 14 kb viral fragment XhoI G (Table 1). The second fragment was amplified from pEcoRI_ASalxh.pBSKS.rev and encoded one of the following regions of *arif-1* (listed as amino acid numbers): K219-D417, T274-D417, P303-D417, or Y320-D417 (Table 1). The resulting plasmids encode ARIF-1 and its truncations fused to the C-terminus of GP64.

To express ARIF-1 with point mutations of proline and tyrosine residues, PCR site-directed mutagenesis was done by amplification of pACT ARIF-1 M1-D417 (full length) using primers to incorporate the desired mutation. Overlapping primers were used to generate proline to alanine mutations at ARIF-1 amino acids P303, P305, P309, P312, P328, P330, P335, P343, P351, P352, and P354 (Table 1), as well as tyrosine to phenylalanine mutations in ARIF-1 at amino acids Y226, Y238, Y241, Y246, Y320, Y325, and Y332 (Table 1). In all cases, the PCR product was purified by agarose gel electrophoresis, digested with DpnI (New England Biolabs) to remove template DNA, transformed into XL-1 Blue *E.coli* (University of California BerkeleyQB3 Macro Lab), and plated on LB agar plates with 100 μg/ml ampicillin (Gibco from Thermo Fisher Scientific). Plasmid DNA from resulting colonies was sequence-verified to ensure the desired changes had been made.

In all cases, to generate DNA ready for transfection, plasmids were transformed into JM109 *E.coli* and cultures grown in 150 ml 2x YT media (MilliporeSigma) overnight at 37°C. A Genelute endotoxin-free Maxiprep kit (MilliporeSigma) was used to purify the DNA.

### Plasmid construction for expression of GFP-tagged cortactin and Arp2/3 complex

To amplify the *S. frugiperda* cortactin gene, total mRNA was isolated from Sf21 cells using an RNeasy kit (Qiagen, Hilden, Germany), and reverse-transcribed to cDNA using a Protoscript II First Strand DNA Synthesis kit (New England Biolabs) using random hexamers as primers. *S. frugiperda* cortactin-specific primers (Table 1) were used to PCR-amplify a 1.9 kb fragment from cDNA that was then used as a template for PCR amplification with a second primer set (Table 1). Next, we constructed a plasmid vector for an N-terminal GFP-tagged *S. frugiperda* cortactin. *eGFP* was PCR-amplified from pEGFPN1 (Takara Bio USA) and inserted using Gibson assembly into a NotI/BamHI-digested pACT (Table 1). The amplified cortactin fragment was then subcloned into the NotI site of the resulting plasmid using Gibson assembly (Table 1). The resulting plasmid expresses GFP fused to the N-terminus of *S. frugiperda* cortactin (GFP-cortactin).

To express a fusion of EGFP to the C-terminus of the p21 (ARPC3) subunit of the Arp2/3 complex (p21*-*EGFP), the *Trichoplusia ni arpc3* gene from pIZ-p21-EYFP (Goley et al., 2006) was PCR-amplified and subcloned using Gibson assembly, along with *eGFP* PCR-amplified from pEGFP-N1 (Takara Bio USA), into the BamHI site of pACT (Table 1).

In all cases, to generate DNA ready for transfection, plasmids were transformed into JM109 *E.coli* and cultures grown in 150 ml 2x YT media (MilliporeSigma) overnight at 37°C. A GenElute endotoxin-free Maxiprep kit (MilliporeSigma) was used to purify the DNA.

### ARIF-1 purification, anti-ARIF-1 antibody generation, and Western blotting

To express recombinant ARIF-1 protein in *E. coli*, the portion of the *arif-1* gene encoding the cytoplasmic C-terminal region (base pairs 654-1254, encoding the C-terminal 199 amino acids) was amplified by PCR from pEcoRI_ASalxh.pBSKS.rev and subcloned into the SspI site of pET-1M (University of California Berkeley QB3 Macro Lab) using Gibson Cloning. This generated the plasmid pET-M1 ARIF-1 219 (Table 1) encoding a fusion protein of the predicted *arif-1* C-terminal cytoplasmic region with an N-terminal 6xHis tag, maltose binding protein (MBP), and tobacco-etch virus (TEV) protease cleavage site (6xHi-MBP-TEV-ARIF-1-219-417). This plasmid was transformed into *E. coli* strain BL21(DE3) (New England Biolabs), the bacteria were grown at 37°C to an OD_600_ of 0.5, and expression was induced with 250 μM IPTG for 2 h. Bacteria were harvested by centrifugation at 4000 rpm for 25 min at 4°C in a Beckman J6M clinical centrifuge (Beckman Coulter Diagnostics; Brea, California), and re-suspended on ice in lysis buffer (50 mM Tris pH 7.5, 200 mM KCl, 1 mM EDTA, 1 μg/mL each leupeptin, pepstatin, and chymostatin (LPC, MilliporeSigma), 1 μg/mL aprotinin (MP Biomedicals LLC, Irvine, CA), and 1 mM phenylmethylsulfonyl fluoride (PMSF, MilliporeSigma)). Lysozyme (MilliporeSigma) was added to the cells at 1 mg/ml; the bacteria were sonicated on ice at 30% power for 4×15 sec in a Branson 450 Digital sonifier and centrifuged at 20,000 × g for 25 min using an SS34 rotor in a Sorvall RC 6+ centrifuge. The supernatant was dripped twice through a 10 ml packed volume of amylose resin (New England Biolabs), washed with 5 column volumes of column buffer (20 mM Tris, pH 7.0, 200 mM NaCl), and eluted with column buffer containing 10 mM maltose. Fractions containing protein were pooled, and protein concentration was determined by Bradford protein assay (Bio-Rad Laboratories, Hercules, CA).

Purified 6xHi-MBP-TEV-ARIF-1-219-417 was used to immunize rabbits (Pocono Rabbit Farm and Laboratory, Canadensis PA) using a 91-day protocol. Before affinity-purifying anti-ARIF-1 antibody, serum was first depleted of anti-MBP antibodies. Buffer exchange was carried out on 10 mg 6xHi-MBP-TEV, purified as described above using an Amicon Ultracell 10 kDa spin concentrator (MilliporeSigma) to concentrate the protein to 20 mg/ml in coupling buffer (0.2M NaHCO_3_, 500 mM NaCl, pH 8.0). This protein was coupled to a 1 ml packed column volume of NHS-activated Sepharose 4 Fast Flow resin (GE Healthcare Life Sciences, Marlborough, MA). 10 ml of serum was diluted 1:1 in binding buffer (20mM Tris, pH 8.0), passed through a 0.22 μm filter, passed over the MBP affinity resin six times at room temperature, and the flow-through was collected. Buffer exchange was carried out as described above to concentrate 10 mg of purified 6xHi-MBP-TEV-ARIF-1-219-417 to 10 mg/mL in coupling buffer. This protein was coupled to another 1 ml packed column volume of NHS-activated Sepharose 4 fastflow resin, and 10 ml of MBP antibody-depleted serum was passed over the 6xHi-MBP-TEV-ARIF-1-219-417 affinity resin. Antibodies were eluted with 100 mM glycine, pH 2.5, and immediately brought to pH 7.5 by addition of 1 M Tris, pH 8.8. Purified antibody was stored at - 20°C.

To observe ARIF-1 expression over the course of early viral infection, Sf21 cells were infected with AcMNPV WOBpos, *AcΔarif-1*, and *AcΔarif-1* rescue viruses at an MOI of 10 and harvested at 0, 2, 4, 6, 8, 10, 12, 16, 20, 24, 28, and 36 hpi. Cells were lysed in protein sample buffer (50 mM Tris, pH 6.8, 10 mM SDS, 370 μM bromophenol blue, 5% glycerol, 1 μg/ml LPC (MilliporeSigma), 1 μg/ml aprotinin (MP Biomedicals LLC), 1 mM PMSF (MilliporeSigma)), and boiled for 5 min. Cell lysates were subjected to SDS-PAGE, transferred to PVDF membrane (Immobilon from MilliporeSigma, Burlington, MA), and probed by Western blotting with rabbit anti-ARIF-1 and rabbit anti-cofilin loading control (provided by Kris Gunsalus; NYU-AD and Michael Goldberg; Cornell).

To observe expression of ARIF-1 and its truncated and mutated derivatives, adherent Sf21 cells were transfected with plasmids expressing ARIF-1 using TransIT-Insect transfection reagent (Mirus Bio, Madison, WI). At 3 d post-transfection, cells were collected, lysed in protein sample buffer (0.2M Tris HCl, 0.4 M DTT, 277 mM SDS, 6mM Bromophenol Blue, 4.3M glycerol), and boiled and subjected to SDS-PAGE and Western blotting as described above.

### Fluorescence microscopy

To image actin structures in live infected cells, Sf21 cells were plated onto 35 mm dishes with 2 mm diameter No 1.5 glass coverslip dishes (MatTek, Ashland, MA) and incubated overnight at 28°C in Grace’s with 10% FBS (Gemini Bio-Products). Cells were transfected with 5 μg of pACT-GFP-actin using TransIT-Insect transfection reagent (Mirus Bio) and incubated for 2 d at 28°C in Grace’s media with 10% FBS and antibiotics (100 μg/ml penicillin/streptomycin and 0.25 μg/ml Amphotericin B). Cells were infected with virus at an MOI of 10, and after 1 h adsorption at 28°C, they were washed with Grace’s media with 10% FBS (this point is defined as 0 hpi), and were incubated at 28°C in Grace’s media with 10% FBS and antibiotics/antimycotics until imaging.

To image actin structures in live cells, Sf21 cells were plated as described above and co-transfected with 5 μg of pACT-ARIF-1 or its truncated or mutated derivatives, pACT-GFP-ARIF-1 (Goley et al., 2006), pACT-GFP-P21, or pACT-GFP-Cortactin. Cells were incubated for 2 d at 28°C in Grace’s media with 10% FBS and antibiotics/antimycotics and imaged.

To quantify formation of clusters of invadosome-like structures in Sf21 cells transfected with pACT-ARIF-1, without or with truncations and mutations, cells were transfected as described above. At 2 d post-transfection, 60 random cells per condition expressing visible GFP-actin were imaged in triplicate at one Z plane at the basal side of the cell. The number of cells with invadosome-like structures, number of invadosome-like structures in each cell, and the shape of the invadosome-like structures were recorded. The data is a result of three biological replicates.

Imaging was performed using a Nikon/Andor confocal microscope with a Yokogawa CSU-XI spinning disc, a Clara Interline CCD camera (Oxford Instruments Inc, Pleasanton, CA), and MetaMorph software (Molecular Devices LLC, San Jose, CA) using a 100X VC objective at 488 nm.

TIRF imaging was performed on a Leica DMi8 S Infinity TIRF HP system with a 100X/1.47 TIRF oil immersion objective, a 488 nm excitation laser, and detected with a Hamamatsu Flash V.4.0 sCMOS camera. Images were processed using ImageJ software.

To quantify invadosome-like structure formation in infected cells over a time course, Sf21 cells were plated onto μclear CELLSTAR black-walled 96-well plates (Greiner Bio-One, Kremsmunster, Austria) and infected at an MOI of 10 with WOBpos, *AcΔarif-1*, or *AcΔarif-1 rescue* virus as described above. Cells were fixed with 4% paraformaldehyde in PHEM buffer (60 mM PIPES, pH6.9, 25 mM HEPES, 10 mM EGTA, 2 mM MgCl_2_), quenched with 0.1 M glycine in PHEM buffer, permeabilized in 0.15% Triton X-100 in PHEM buffer, and blocked with 5% normal goat serum (MP Biomedicals, Irvine, CA) and 1% bovine serum albumin in PHEM. Cells were stained with anti-GP64 B12D5 primary antibody (a gift from Dr. Loy Volkman) at a 1:200 dilution in PHEM buffer and with a secondary goat anti-mouse AlexaFluor 488 conjugated antibody (Invitrogen from Thermo Fisher Scientific) at a 1:400 dilution, both in PHEM buffer. F-actin was visualized with Alexa Fluor 568 Phalloidin (Invitrogen from Thermo Fisher Scientific) diluted 1:200 in PHEM buffer, and DNA was visualized with 5 μg/ml Hoechst (MilliporeSigma) in PHEM buffer. Cells were imaged with an Opera Phenix high-content image screening system (PerkinElmer, Waltham, MA) using a 40x water immersion objective (PerkinElmer). Images were analyzed on Harmony image analysis software (PerkinElmer) using maximum intensity projections, and the number of cells, number of cells with GP64 signal, and number of cells with clusters of invadosome-like structures, and the number of clusters of invadosome-like structures in each cell were recorded. The data is a result of three biological replicates.

For Arp2/3 complex drug inhibition experiments, Sf21 cells were plated and transfected with pACT GFP-actin as described above. Cells were infected with WOBpos virus at MOI of 10 as described above, and at 4 hpi, latrunculin A in DMSO was added to a final concentration of 4 μM, or CK666 or CK689 in DMSO was added to a final concentration of 100 μM. Imaging was begun immediately, with 8 cells imaged every 30 s. Images were processed in image J, and the percent of invadosome-like actin structures remaining was recorded. The data is a result of three biological replicates.

## Acknowledgements

We wish to thank the following individuals for their contributions to this work: Loy Volkman for her advice, support, and role as a pioneer in the field of baculovirus host-pathogen interactions; Mary West and Chris Noel of the UC Berkeley QB3 High-Throughput Screening facility for use of facilities and technical support; Holly Aaron and Jen-Yi Lee of the UC Berkeley Molecular Imaging Center for support for our microscopy work; Michael Goldberg and Kris Gunsalus for the rabbit anti-cofilin GA15 antibody; Susan Hepp and Bisco Hill for providing plasmid constructs and advice. This work was supported by grant 1S10OD021828-01A1 from the NIH Office of the Director, which funded the Opera Phenix microscope; and grant R35 GM127108 from the NIH/NIGMS to M.D.W.

