## Supplementary figures and Video Legends for "Baculovirus actin-rearrangement-inducing factor ARIF-1 induces the formation of dynamic clusters of invadosome-like structures"

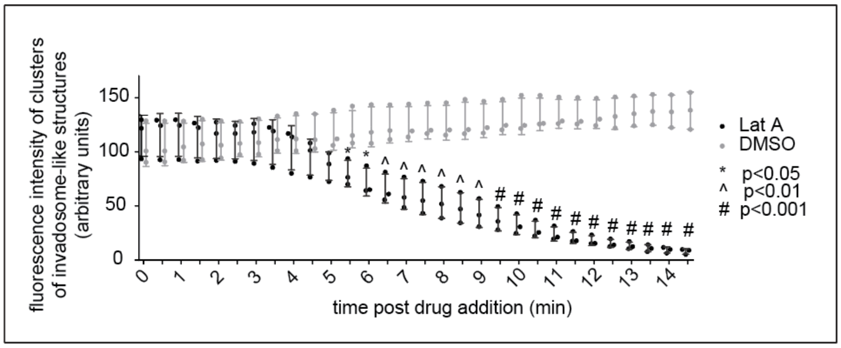


**Supplemental Figure S1: Latrunculin A induces dissipation of clusters of invadosome-like structures.** Individual clusters of invadosome-like structures were identified in live Sf21 cells transiently expressing GFP-actin and infected with AcMNPV MOI of 10, 2 d post-transfection. 4 μM Latrunculin A or DMSO was added to cells, and fluorescence of clusters of invadosome-like structures was measured from 0 to 14.5 min post-drug addition. Data are mean ± SD of three biological replicates of 38 individual invadosome clusters. P-values were calculated using multiple t-tests and are indicated as follows: * = $p<0.05$, ^ = $p<0.01$, # = $p<0.001$.

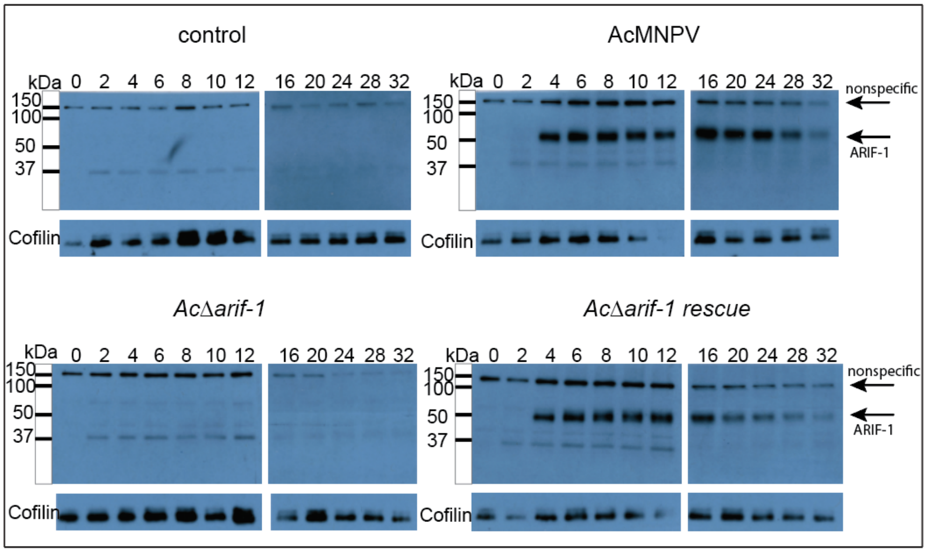


**Supplemental Figure S2: The timing of ARIF-1 expression corresponds with formation of clusters of invadosome-like structures.** Western blots of lysates of Sf21 cells infected with the indicated virus harvested at 0 to 32 hpi. Lysates were probed with a rabbit anti-ARIF-1 antibody. Nonspecific and ARIF-1 bands are indicated by arrows. Cofilin is shown as a loading control.

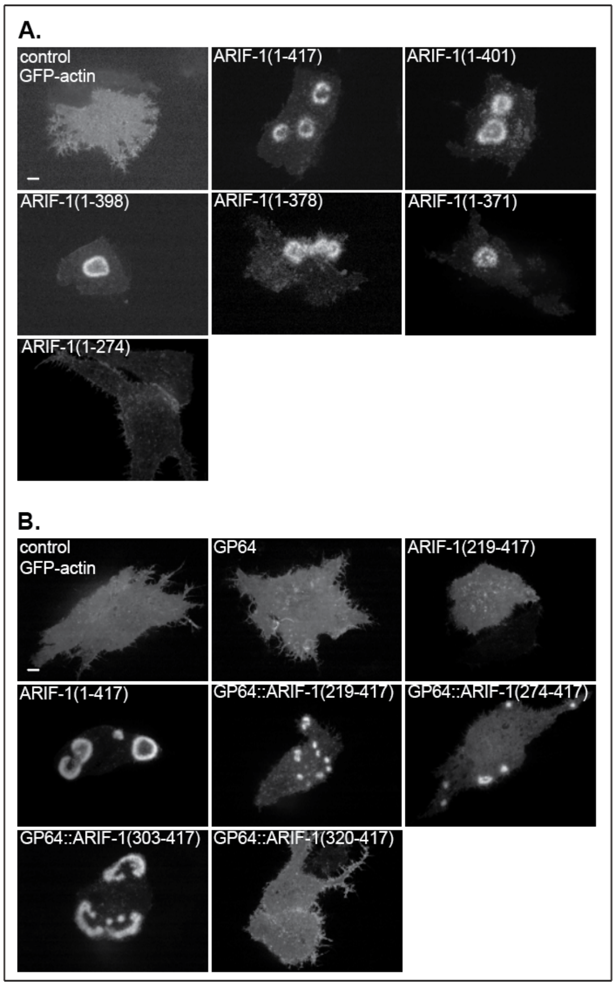


**Supplemental Figure S3: Cells expressing the ARIF-1 C-terminal cytoplasmic region form clusters of invadopodia-like structures. (A)** Confocal images of Sf21 cells transiently expressing GFP-actin and ARIF-1 C-terminal truncations. Images were taken 2 d post-transfection and are representative of three biological replicates. Scale bars = 5 µm. **(B)** Confocal images of Sf21 cells transiently expressing GFP-actin and ARIF-1 N-terminal truncations and GP64::ARIF-1 truncations. Images were taken 2 d post-transfection and are representative of three biological replicates. Scale bars = 5 µm.

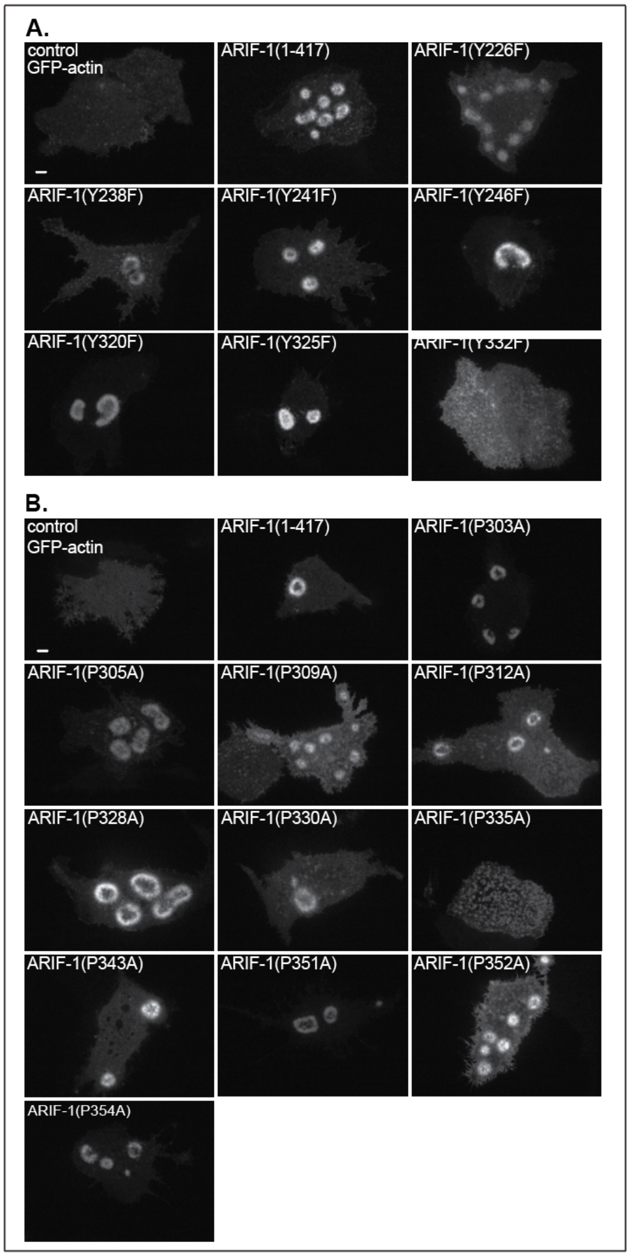


**Supplementary Figure S4: Invadosome-like structures form in cells expressing ARIF-1 tyrosine and proline residue mutations. (A)** Confocal images of Sf21 cells transiently expressing GFP-actin and the indicated tyrosine residue ARIF-1 point mutations. Images were taken 2 d post-transfection and are representative of three biological replicates. Scale bars = 5 µm. **(B)** Confocal images of Sf21 cells transiently expressing GFP-actin and ARIF-1 with the indicated proline residue point mutations. Images were taken 2 d post-transfection and are representative of three biological replicates. Scale bars = 5 µm.

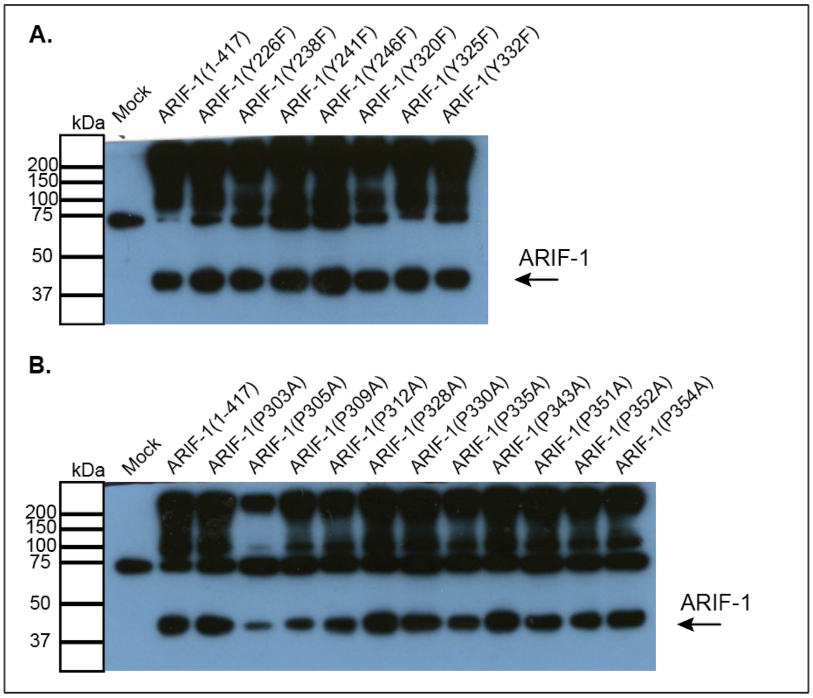


**Supplementary Figure S5: ARIF-1 is expressed in cells transiently expressing ARIF-1 proline and tyrosine residue point mutants. (A)** Western blot of lysates of Sf21 cells transiently transfected with ARIF-1 tyrosine residue point mutants. Blots were probed with rabbit anti-ARIF-1 antibody. ARIF-1 signal is indicated by the arrow. **(B)** Western blot of lysates of Sf21 cells transiently transfected with ARIF-1 proline residue point mutants. Blots were probed with rabbit anti-ARIF-1 antibody. ARIF-1 signal is indicated by the arrow.

**Supplementary Video Legends**

**Video S1: Dynamic invadosome-like structures form in AcMNPV infected Sf21 cells.** Confocal timelapse imaging of Sf21 cells expressing GFP-actin and infected with AcMNPV. Cells were imaged every 2 min starting at 140 min post infection. Video is 151 frames shown at 5 frames per s. Scale bar is 5 µm. Time post AcMNPV infection is shown.

**Video S2: Stationary invadosome-like structures form clusters.** Confocal timelapse imaging of Sf21 cells expressing GFP-actin and infected with AcMNPV. Cells were imaged every 2 min starting at 218 min post infection. Video is 13 frames shown at 5 frames per s. Red circles indicate individual invadosome-like structures which disappear, modulating the shape of the larger cluster. Yellow circles indicate individual invadosome-like structures that are maintained. Scale bar is 5 µm. Time post AcMNPV infection is shown.

**Video S3: ARIF-1 expression is sufficient for formation of dynamic invadosome-like structures.** Confocal timelapse imaging of Sf21 cells expressing GFP-actin and ARIF-1. Cells were imaged every 30 s starting 48 h post transfection. Video is 30 frames shown at 5 frames per s. Scale bar is 5 µm. Time elapsed during imaging is shown.

**Video S4: Plasma membrane targeted ARIF-1 C-terminal cytoplasmic region is sufficient for formation of dynamic clusters of invadosome-like structures.** Confocal timelapse imaging of Sf21 cells expressing GFP-actin and GP64::ARIF-1(219-417). Cells were imaged every 30 s starting 48 h post transfection. Video is 91 frames shown at 5 frames per s. Scale bar is 5 µm. Time elapsed during imaging is shown.

**Video S5: Invadosome-like structures are uniformly distributed along the basal PM in Sf21 cells expressing ARIF-1(Y332F).** Confocal timelapse imaging of Sf21 cells expressing GFP-actin and ARIF-1(Y332F). Cells were imaged every 30 s starting 48 h post transfection. Video is 121 frames shown at 5 frames per s. Scale bar is 5 µm. Time elapsed during imaging is shown.

**Video S6: Invadosome-like structures are uniformly distributed along the basal PM in Sf21 cells expressing ARIF-1(P335A).** Confocal timelapse imaging of Sf21 cells expressing GFP-actin and ARIF-1(P335A). Cells were imaged every 1 min starting 48 h post transfection. Video is 81 frames shown at 5 frames per s. Scale bar is 5 µm. Time elapsed during imaging is shown.

**Video S7: Latrunculin A induces rapid dissolution of clusters of invadosome-like structures.** Confocal timelapse imaging of Sf21 cells expressing GFP-actin and infected with AcMNPV. At 4 hpi, 4 µM Latrunculin A was added to cell medium and cells were imaged every 1 min for 1.35 h. Video is 81 frames shown at 5 frames per s. Scale bar is 5 µm. Time elapsed after viral infection and Latrunculin A addition is shown.

**Video S8: Arp2/3 complex inhibitor CK666 induces dissolution of clusters of invadosome-like structures.** Confocal timelapse imaging of Sf21 cells expressing GFP-actin and infected with AcMNPV. At 4 hpi, 100 μM of CK666 was added to cell medium and cells were imaged every 30 s for 36.5 min. Video is 73 frames shown at 5 frames per s. Scale bar is 5 μm. Time elapsed after viral infection and CK666 addition is shown.

**Video S9: Inactive control Arp2/3 inhibitor CK869 has no effect on clusters of invadosome-like structures.** Confocal timelapse imaging of Sf21 cells expressing GFP-actin and infected with AcMNPV. At 4 hpi, 100 μM of CK869 was added to cell medium and cells were imaged every 30 s for 66.5 min. Video is 133 frames shown at 5 frames per s. Scale bar is 5 μm. Time elapsed after viral infection and CK666 addition is shown.
